# The olfactory bulb carries out concentration invariance calculations by itself and does it very quickly

**DOI:** 10.1101/2022.08.17.504274

**Authors:** Yunsook Choi, Lawrence B. Cohen

## Abstract

The olfactory bulb is known to carry out computations for the olfactory perception of concentration invariance of odor recognition, a perception that is essential for life as we know it. However it is not known whether these computations require the extensive feedback to the bulb from the many brain regions which send this feedback. We have measured the concentration dependence of the mitral cell output of the bulb before and after blocking this feedback by lidocaine infusion into the medial olfactory peduncle. Surprisingly we found that the concentration dependence of the mitral cell output was unaffected by the lidocaine block of feedback. Furthermore the computation is rapid; it is complete within a single 250 msec inhalation. The computations for concentration invariance are most likely carried out within the olfactory bulb itself and they are carried out quickly.

## INTRODUCTION

The olfactory bulb is known to carry out computations for the perception of concentration invariance of odor recognition (Storace and Cohen, 2017; Bolding and Franks 2018). Without this perception it would not be possible to follow odor concentration gradients to locate objects of critical importance such as food, mates, prey, and predators. However, the experiments reporting the bulb computations about concentration invariance did not address the question of whether the computation was carried out by the bulb itself or in collaboration with feedback to the bulb from higher brain regions.

This feedback has been known anatomically for more than a century (Ramon y Cajal, 1904; Haberly and Price, 1978:I; Haberly and Price, 1978:II; Luskin and Price, 1982; Kerr and Hagbarth,1955; Macrides et al., 1981; Mclean and Shipley, 1987; reviewed in Shipley and Ennis, 1996) and includes modulatory feedback from the basal forebrain, raphe nuclei, hypothalamus, and locus coeruleus as well as feedback from both olfactory and non-olfactory cortex. The feedback is “very dense” (Shipley and Ennis, 1996) with more feedback axons than feedforward axons from the olfactory epithelium.

### Previous results

The feedback from higher centers has been shown to be important for a number of olfactory responses. Some of these occur over long time scales which could involve modulatory feedback. Some examples follow. *Blocking feedback*: Mandairon et al., (2014) found that visual context dependent olfactory behavior was blocked by lidocaine infusion into the olfactory peduncle. In addition, the spatial expression of the early activity gene, Zif268, was altered by lidocaine infusion into the peduncle. Martin et al., (2006) found that lidocaine infusion into the peduncle greatly reduced the bulb β-oscillations in response to odorants while increasing the γ-oscillations. These changes occurred only in trained animals. Kiselycznyk et al., (2006) found that electrical lesions to the olfactory peduncle blocked olfactory learning about food rewards. Finally, Otasu et al., (2015) found that inactivation of piriform cortex increased odor responsiveness and pairwise similarity of mitral cells.

### Activating feedback

Medinaceli Quintela et al., (2020) found that activating neurons in the anterior olfactory nucleus would dramatically inhibit the mitral cell response to odorant. Eckmeier and Shea, (2014) found that activating the locus coeruleus would inhibit the input to the bulb from the olfactory receptor cells in the olfactory epithelium. Because of these and several other similar observations, it would be reasonable to hypothesize that feedback from higher brain centers would also be important for the computation of concentration invariance. The results in this paper provide a test of this hypothesis. For reviews about feedback to the olfactory bulb see Mandairon and Linster (2009); Restrepo et al., (2009); Brunert and Rothermel (2021).

### Method

A procedure for blocking the feedback to the bulb was first developed by C.M. Gray and J.E. Skinner in 1986. They reported that simultaneous cooling in the region of the olfactory peduncle and the lateral olfactory tract would block >95% of the olfactory bulb response to stimulation of both the anterior commissure and the lateral olfactory tract. This blockade increased olfactory bulb local field potential oscillations (Grey and Skinner, 1986). However there was noticeable crosstalk and cooling one site affected the bulb response to stimulation of the other site. Martin et al., (2006) developed a feedback only blockade by injecting lidocaine into the medial olfactory peduncle. This infusion increased the γ-oscillations in the bulb without effecting the bulb response to stimulation of the lateral olfactory tract. We have used the procedure developed by Martin et al., (2006) to study the effect of blocking feedback on the concentration invariance computation.

To determine the effect of feedback on the input-output transformation for concentration invariance we have measured the effect of lidocaine infusion into the medial olfactory peduncle on the concentration dependence of the mitral/tufted cell output activity. The mitral/tufted cell activity was measured with the calcium sensor, GCaMP6f that was expressed via infusion of AAV-floxed GCaMP6f into the olfactory bulbs of Tbx21-Cre mice (Nagai et al., 2005) that express cre recombinase specifically in the mitral/tufted cells. The GCaMP6f fluorescence from olfactory bulb glomeruli was measured with wide-field microscopy rather than 2-photon microscopy because wide-field microscopy had a larger field of view which allows recording from a larger number of glomeruli.

## METHODS

### Animals

All animal experiments were performed in compliance with institutional guidelines and protocols approved by the Institutional Animal Care and Use Committee at the Korea Institute of Science and Technology. Male and female adult mice (10-20 weeks old) were used. Tbx21-Cre transgenic mice used in this study were acquired from the Jason Laboratory (Jax #024507). Tbx21-cre mice express *cre* recombinase in mitral/tufted cells of the olfactory bulbs (Nagai et al., 2005; Chen et al., 2013).

### Surgical procedures

#### Mice preparation for optical imaging

Mice were anesthetized with Ketamine/Xylazine (initial dose 90mg/10mg/kg body weight). Anesthesia was maintained by additional injection of Ketamine/Xylazine (45mg/5mg/kg) and anesthesia depth was monitored using the toe pinch reflex. The animal body temperature was maintained at ~36.5°C using a homeothermic heating pad). The animal breathed freely throughout the experiment. Mice were kept hydrated by intraperitoneal injections of Ringer’s solution.

The craniotomy over the olfactory bulb was carried out as described previously (Braubach et al., 2018). Local anesthetic (1% Bupivacaine) was applied at the incision sites. A longitudinal incision was made from behind the ear to the anterior part of the olfactory bulb. The skin was retracted to expose the skull. A head-post was attached to the posterior skull with cyanoacrylic glue and dental cement (Vertex Orthoplast, Vertex-Dental, Zeist, Holland). The skull above the dorsal olfactory bulb was thinned with a dental drill (Foredom K.1070), and the craniotomy (~1.5 × 2 mm) was performed by removing the thinned bone. 2% agarose in Ringer’s solution and then a glass window were placed over the olfactory bulb.

#### Virus injection

AAV-GCaMP6f (AAV1.Syn.Flex.GCaMP6f.WPRE.SV40, Lot #CS1192-NZ) were purchased from the Penn Vector Core. The virus was injected into Tbx21-cre transgenic mice near the mitral cell layer (Figure 1). Mice were anesthetized and a hole (0.5 mm) was drilled using a stereotaxic instrument (Neurostar, Germany) in the same hemisphere where the guide cannula was implanted into the olfactory peduncle. GC (genome copy/ml) titer for Adeno-associated viral (AAV) vectors was 4.38e13 and was diluted in sterile saline by 4 fold. Virus was withdrawn into the 10 uL syringe (World Precision Instruments, Sarasota, USA) and 1uL of the diluted virus was injected into the bulb using 35 gage metal needle (World Precision Instruments). Virus was injected 450 um below the surface of the bulb (Figure 12) at the rate of 50nL/min using a stereotaxic (Neurostar, Germany). Mice were given acetaminophen-treated water (1.6mg/ml) for 5 days after virus injection. The optical imaging was carried out between 14 and 28 days after virus injection to allow for the spread of the AAV-GCaMP6f virus and the GCaMP6f expression.

**Figure 1.**
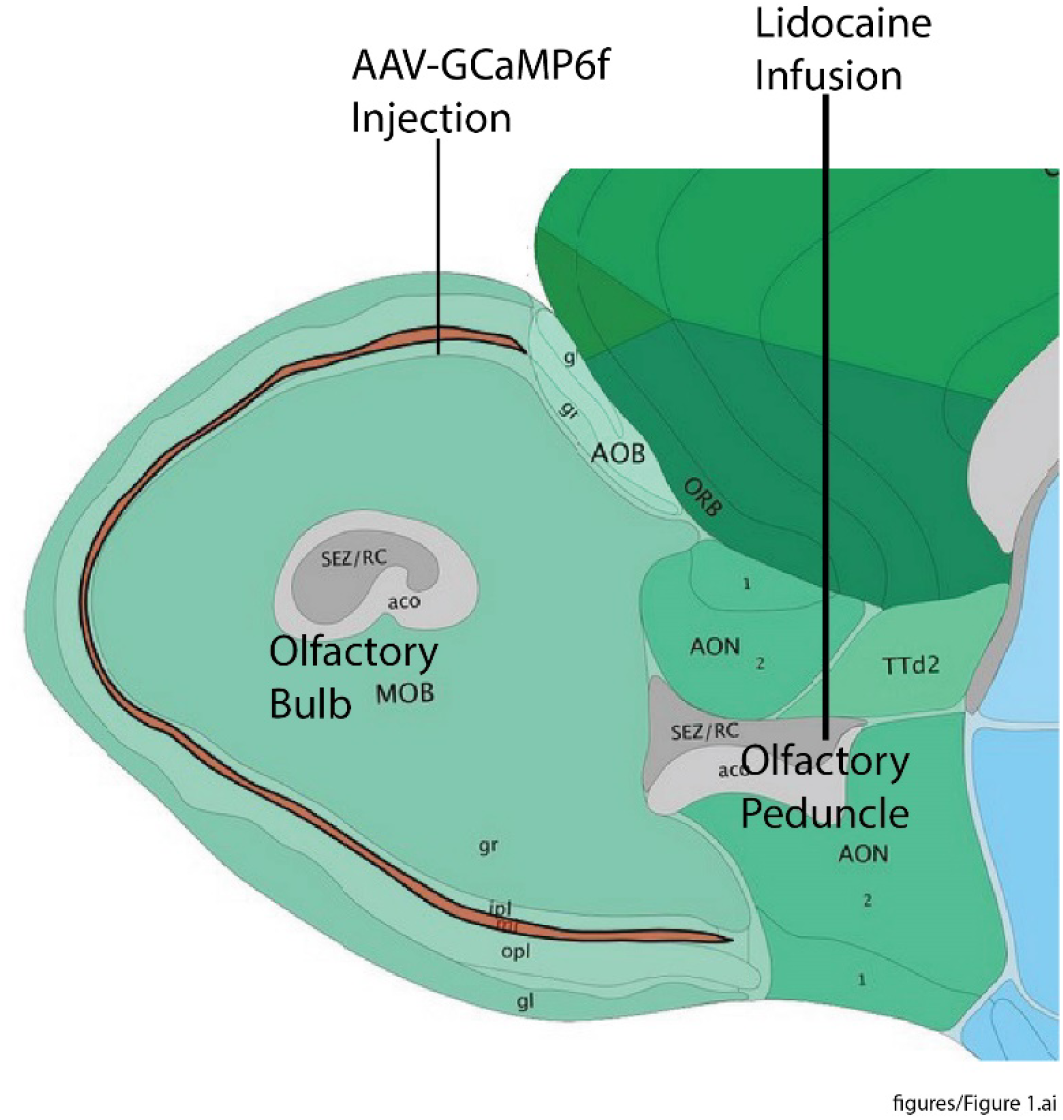
Schematic diagram: AAV-GcaMP6f injection and lidocaine infusion.

#### Lidocaine cannula implantation

Mice were anesthetized with a mixture of ketamine (100 mg/kg) and xylazine (10 mg/kg) and placed in a stereotaxic instrument (Neurostar, Tubingen, Germany). A hole (0.6 mm) was made in the skull for the guide cannula implantation. Mice were implanted unilaterally with a guide cannula (26 gage, PlasticsOne, Roanoke, USA) into the medial olfactory peduncle at the following coordinates with respect to the bregma: AP, +2.4mm; ML, +0.75 mm; DV, −3 mm (Figure 1). The tip of the guide cannula was positioned 1 mm above the infusion site. A dummy cannula was inserted into the guide cannula to prevent infection and blockage. Two mounting screws drilled into the skull and dental cement were used to help secure the guide cannula to the skull. Mice were allowed to recover for 14 days.

### Infusion procedure

Lidocaine was infused into anesthetized or awake mice. The internal cannula (33 gage, PlasticsOne, Roanoke, USA) was connected to a 5 uL syringe (Hamilton) through a polyethylene tube. The dummy cannula was removed and internal cannula was inserted into the guide cannula. The internal cannula protruded 1 mm beyond the tip of the guide. Lidocaine (Sigma, 2%) was infused into the olfactory peduncle at the rate of 0.25 uL/min using a syringe pump. The total volume was 2 uL. The internal cannula was left in place for 5 min after infusion to prevent backflow. The dummy cannula was then replaced. Then optical imaging or electrode recording started.

### Wide-field fluorescence imaging

Wide-field optical imaging of the dorsal olfactory bulb was done using the wide-field section of a modified Sutter MOM (Sutter instruments, Novato, USA) equipped with a RedShirtImaging NeuroCCD SM-256 camera (RedShirtImaging, Decatur, GA). The light from a Prizmatix LED (UHP-T-LED-460-LN) was used with a Zeiss 5x, 0.25 NA objective. GCaMP6f fluorescence was measured using a 472/30□nm band-pass excitation filter (Semrock FF02-472/30), a 495 nm dichroic mirror, and a 496□nm long pass emission filter (Semrock FF01-496/□P). The emitted fluorescence was recorded at 125 frames/second using NeuroPlex software (RedShirtImaging).

### Mice preparation and recording spontaneous local field potentials

A stainless-steel electrode was screwed in the skull bone directly above the olfactory bulb in the same hemisphere that was implanted with a lidocaine guide cannula. The reference electrodes were positioned in the skull over the cerebellum. Mice recovered for 14 days before recording. The electrodes were connected to a differential AC amplifier (A-M systems model 1700). The output signal from the AC amplifier over the frequency range between 1.0Hz and 500Hz was recorded using a digital data acquisition system (Axons Instruments, Digidata 1440A Digitizer, San Jose, USA). Recording was done in awake mice for 30 min before the lidocaine infusion and continued for ≥30 min after lidocaine infusion. The recording was filtered with a band-pass filter 65-90 Hz using Clampfit software (Axon Instruments, San Jose, USA).

### Imaging data analysis

Imaging data were analyzed with NeuroPlex (RedShirtImaging) software. The location of glomeruli that have calcium signals in the image was determined with the Frame Subtraction function (Wachowiak and Cohen, 2001). The temporal average of the 1-2s preceding the odorant stimulus was subtracted from a 1-2s period during the response to odorants and the software displayed the pixels with fluorescence intensity changes.

The fluorescence signals were converted to fractional fluorescence changes (ΔF/F) by dividing the fluorescence intensity change in the glomerulus by the fluorescence intensity preceding the stimulus. Calcium imaging data were collected either as single trials or the average of 4 trials. Trials were separated by a minimum of 45 sec.

The calcium signal in each glomerulus evoked by 0.45% of saturated vapor was plotted as the fraction of the signal size evoked by 9% (Figures 3 and 4). In most experiments, these calculations were carried out for the same glomeruli before and after lidocaine infusion.

### Odorant delivery

The odorant, methyl valerate (Sigma-Aldrich), was delivered using an olfactometer described previously (Vucinic et al., 2006). The odorant was diluted from saturated vapor with cleaned humidified air.

The figures were made using Excel, Origin, and Adobe Illustrator.

## RESULTS

Two sets of experiments were carried out on separate groups of mice. In the first set we measured the effect of lidocaine infusion into the olfactory peduncle on the γ-oscillations in the olfactory bulb. In the second experiment, carried on different mice, we measured the effect of lidocaine infusion on the odorant concentration dependence of the mitral/tufted cell output of the olfactory bulb using the activity indicator GCaMP6f. We were not able to carry out both experiments on one animal because of the limited space on the bone covering the mouse brain. We used anesthetized mice for these experiments to be consistent with the experiments of Storace and Cohen (2017) which showed that the olfactory bulb made a substantial contribution to the computation converting the input to the bulb into a relatively concentration invariant output.

### Effect of lidocaine block on the γ-oscillations in the olfactory bulb

Following the procedures of (Martin et al., 2006), we implanted a lidocaine infusion cannula and used bone screw electrodes for recording the γ oscillations in the olfactory bulb. We checked the effectiveness of our lidocaine infusions by measuring changes in the γ oscillations in response to the infusion of 2µl of 2% lidocaine into the medial olfactory peduncle. Figure 2 illustrates typical 24 second sections of a 30 minute recording made before and after lidocaine infusion. The recordings were band-pass filtered between 65 to 90 Hz using Clampfit software. The lidocaine infusion dramatically increased the γ oscillations during the 30 minute recording. A similar dramatic increase was observed in three out of three preparations. Thus, we are reasonably confident that our lidocaine infusion procedure blocks a substantial fraction of the feedback to the bulb from higher olfactory centers.

**Figure 2.**
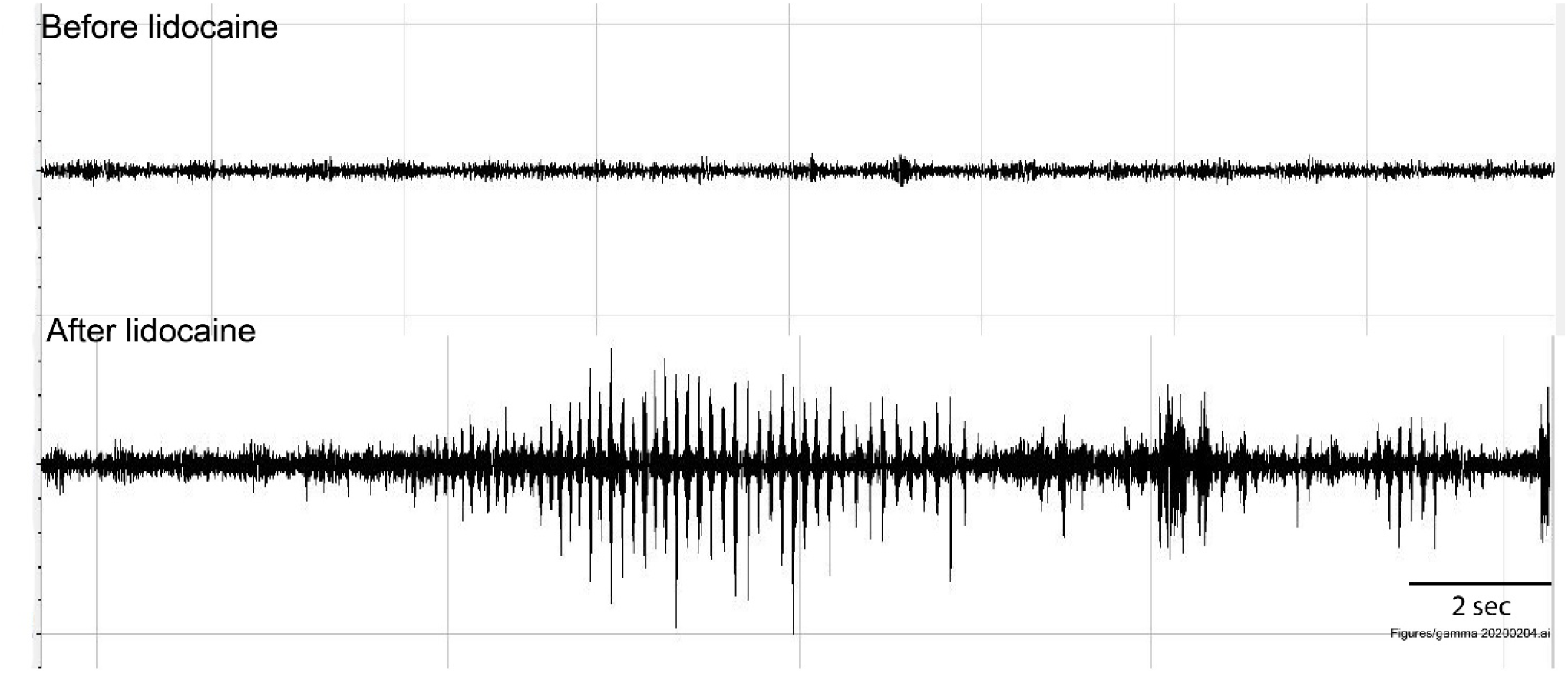
Lidocaine infusion into the olfactory peduncle increases olfactory bulb gamma oscillations before lidocaine

### Effect of lidocaine infusion on the concentration dependence of the mitral/tufted cell activity

We carried out experiments to examine the effect of lidocaine infusion into the peduncle on the mitral/tufted cell output in response to high and low odorant concentrations. These experiments are technically challenging. We start with AAV-GCaMP6f virus injection into the mitral cell layer of Tbx21-Cre mice (Nagai et al., 2005). Following Martin et al, (2006) and Mandairon et al, (2014) mice were then implanted with a guide cannula (26-gauge; Plastics One Roanoke, VA, USA) into the medial olfactory peduncle. Two supporting screws were added. Two weeks later a craniotomy was performed over the olfactory bulb and a head-post glued to the caudal bone. This uses all of the available real estate on the dorsal bone. Finally optical recordings of olfactory responses to odorant presentations were measured using wide-field microscopy. Then lidocaine was infused into the medial peduncle at the rate of 0.25 μl/min. Starting 10 minutes after lidocaine infusion to allow time for lidocaine diffusion, we determined the signal size in the output activity in response to 9% and 0.45% of saturated methyl valerate vapor.

We found that lidocaine injection did not affect the organization of the glomerular map in response to odorant presentation (data not shown). Surprisingly, Figure 3 shows that blocking feedback from higher olfactory centers by lidocaine infusion did not affect the concentration dependence of the olfactory bulb output. For each glomerulus with a clear response to odorant presentations at both high and low concentrations, we set the response to 9% odorant to 1.0 and plotted the size of the response to the low concentration as a percentage of the response to 9%. Each data point in Figure 3 represents the response of one glomerulus; from one to four measurements were made in each condition (before or after lidocaine infusion). If blocking feedback had blocked the computation of the perception of concentration invariance, then we would have expected that the 0.45% response after lidocaine would be much smaller than that before lidocaine. However the result in Figure 3 shows that the response to the low concentration was unaffected by the lidocaine infusion. Figure 4 illustrates the results similar experiments carried out on four additional mice. The average effect of lidocaine in the five experiments was a very small increase in the low concentration response: +0.3% ± 3.2% (S.E.M.). The t test p value for a difference from zero effect was 0.92. The difference between the input and the relatively concentration invariant output in Figure 3 of Storace and Cohen (2017) was ~30%. Thus, lidocaine infusion had no substantial effect on the concentration invariance of the mitral cell output in response to the odorant.

**Figure 3.**
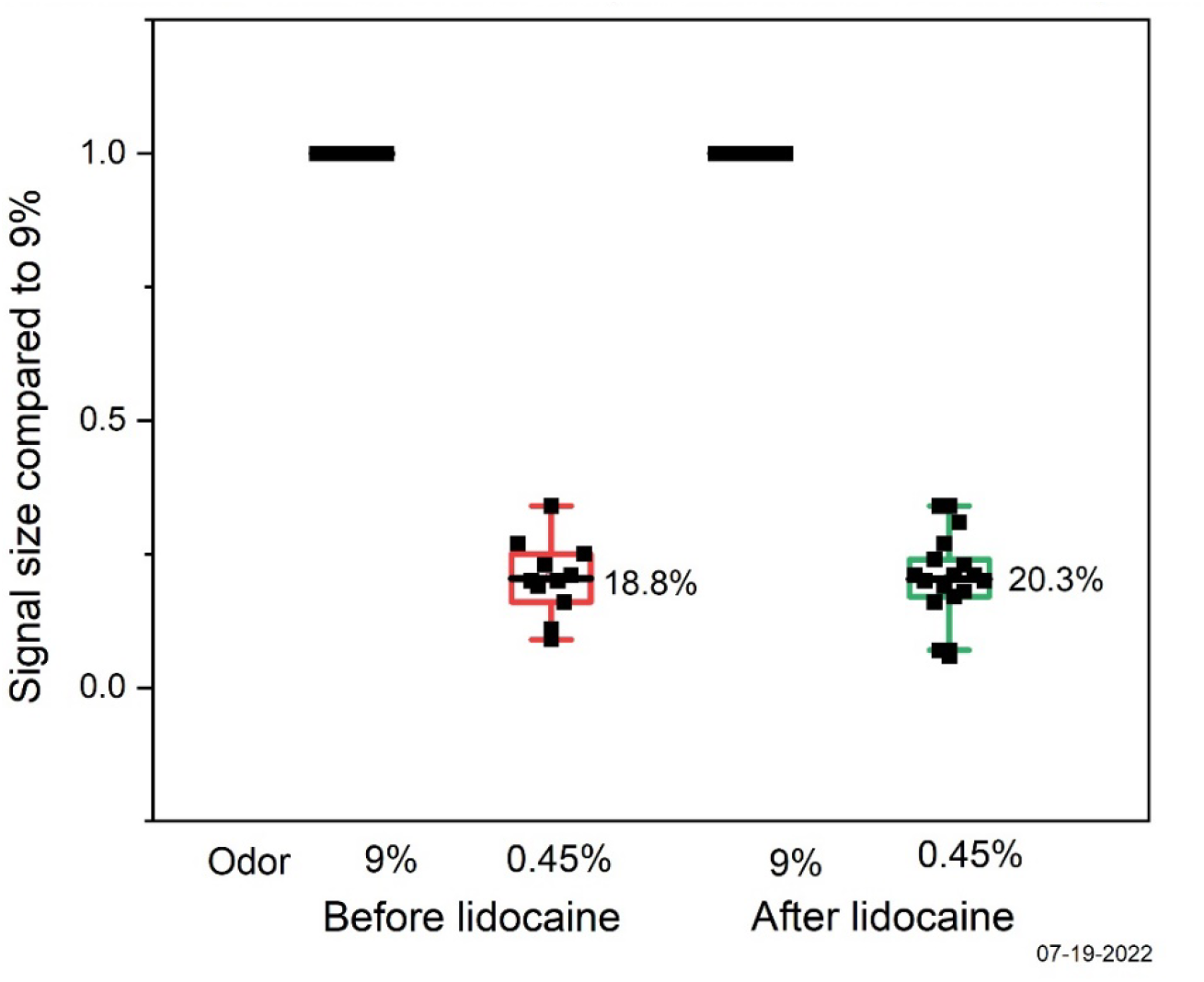
Effect of lidocaine on the concentration dependence of the mitral/tufted response

**Figure 4.**
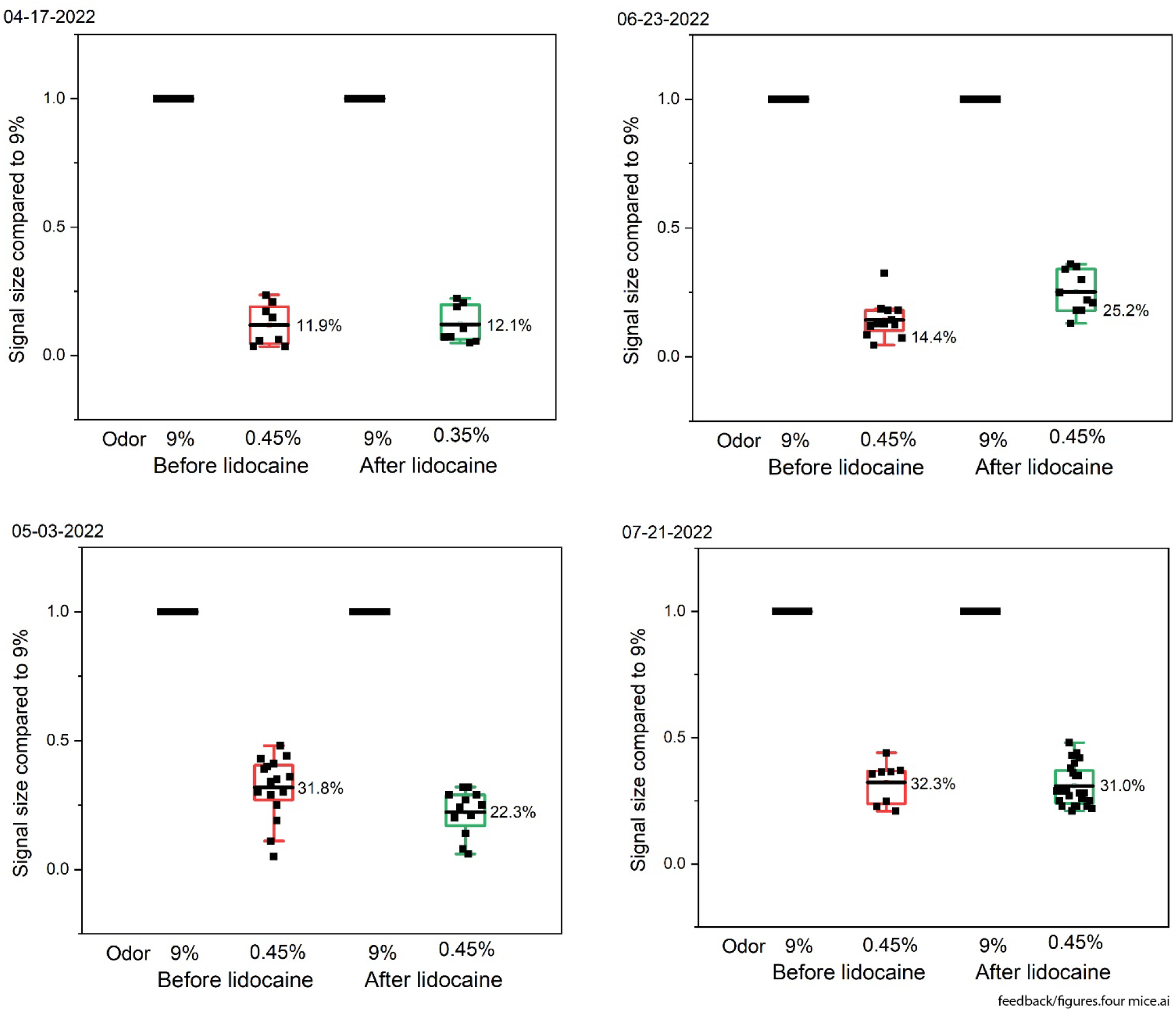
Effect of lidocaine on the mitral/tufted response to odorants

The results in Storace and Cohen (2017) showed that the computation of concentration invariance of the mitral/tufted cell output occurred within a two second odorant presentation. We asked whether the computation might already be achieved in the first breath following an odorant presentation. We compared the concentration dependence of the output from the first inhalation with the output averaged over the two second period. Figure 5 shows that the mitral/tufted output has the same concentration dependence for the first inhalation (red) and the average of the two second odorant exposure (black). Each data point represents the mean output signal from eight glomeruli. The data are taken from the experiment described in Figure 2 of Storace and Cohen (2017). The respiratory frequency was ~2Hz; each inhalation would occupy ~250 msec. Thus, the computation that is responsible for the concentration invariant output must be rapid compared to the ~250 msec time scale of an inhalation.

**Figure 5.**
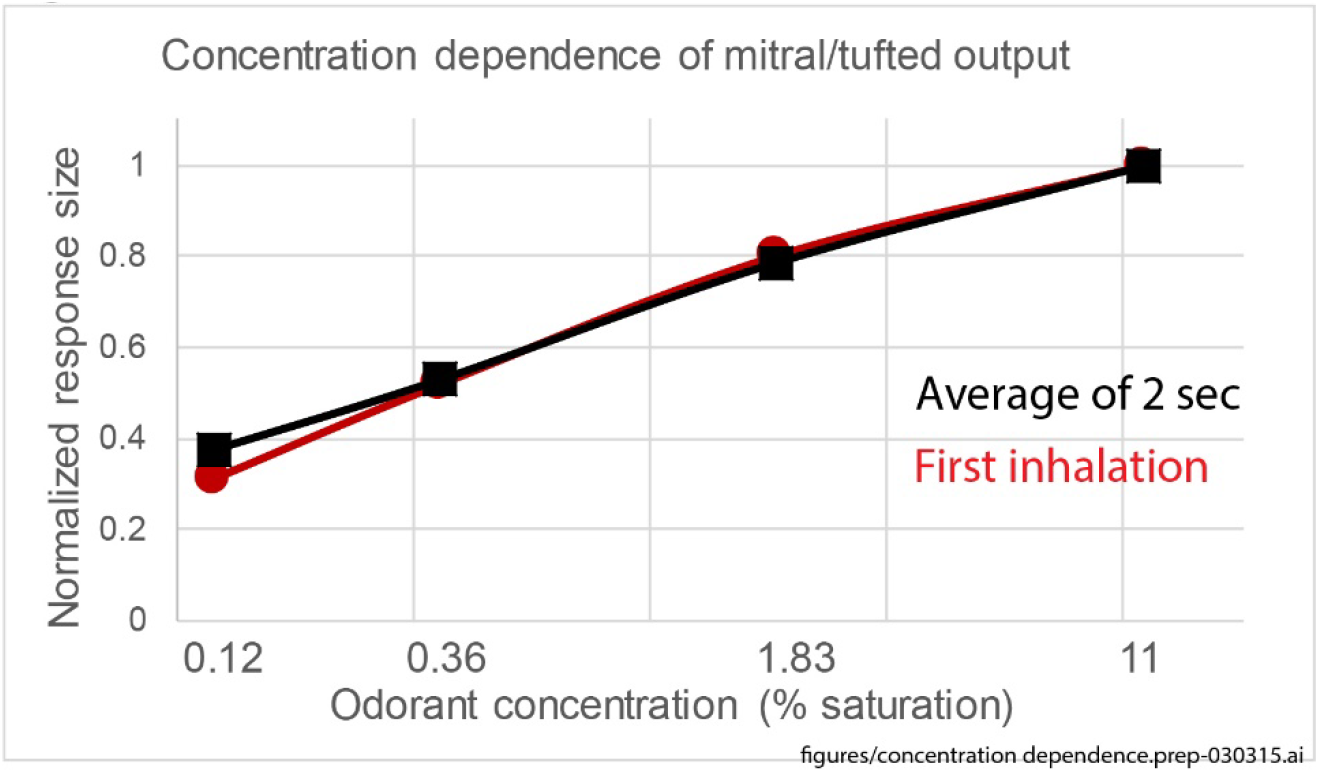

### Effect of lidocaine infusion on the amplitude of the odorant response

Even though the lidocaine infusion did not affect the concentration dependence of the olfactory response, it did result in a substantial reduction in the odorant response size. The mean reduction in the mitral/tufted signal size after lidocaine infusion in the five mice was 56.2 ± 5.3% (S.E.M.). The trial to trial differences where both trials were either before or after lidocaine infusion was much smaller; mean −12.8% ± 10.3% (S.E.M.), n=7 comparisons.

## DISCUSSION

### Effectiveness of lidocaine infusion into the medial olfactory peduncle on blocking feedback

We recorded two dramatic effects of lidocaine infusion: the increase in γ-oscillations recorded in the bulb (Figure 2) and the 50% decrease in the odorant response amplitude of the mitral/tufted cells. Because there was a large effect of the lidocaine infusion, we think that it is likely that a large fraction of the feedback was blocked.

### Effect of lidocaine infusion on the mitral/tufted cell odorant response amplitude

Lidocaine infusion resulted in a 50% decrease in the response amplitude of the mitral tufted cells by the time of our first measurement, 10 minutes after the end of the infusion. This decrease was then relatively stable for at least 30 minutes. However, we don’t know the time course of this lidocaine effect. One hypothesis is that the inhibition results from blocking net excitatory modulatory inputs.

### Effect of lidocaine infusion on the concentration dependence of the mitral/tufted odorant response

We found that lidocaine block of feedback to the olfactory bulb had no detectable effect on the concentration dependence of the output of the bulb carried by the mitral/tufted neurons. Thus the conversion of the bulb input, a confound of odor concentration and odor quality (e.g. Wachowiak and Cohen, 2001), into an output that is relatively concentration invariant (e.g., Storace and Cohen, 2017; Bolding and Franks, 2018), is carried out even when feedback from other brain regions is blocked. This was surprising considering the very dense anatomical feedback to the olfactory bulb and the many reports showing that this feedback affected bulb function.

Because olfaction is used by animals to find odor sources in an environment of turbulent odorant plumes, the computation of concentration invariance must be carried out continuously and very rapidly. The result in Figure 5 showed that the calculation of the concentration invariance is indeed rapid; it is complete during a single inhalation. This computation might be more efficient if it were carried out within the olfactory bulb rather than depending on feedback from higher brain centers.

It may be difficult to determine the actual time dependence of the transformation from a confounded input into a concentration invariant output with currently available tools. Genetically Encoded Calcium Indicators (GECIs) are slow relative the expected time scale and Genetically Encoded Voltage Indicators (GEVIs) have signal-to-noise ratios that are unlikely to provide clear signals in a high time resolution measurement (Storace et al., 2015).

The retina is another brain region which carries out important and rapid transformations of sensory input into more complicated outputs. It carries out these transformations with very little feedback from higher brain centers; in the mouse there are only 150 axons providing feedback from the brain. This represents only 0.2% of the axons in the optic nerve (Gastinger et al., 2006; Repérant et al., 2006). These axons are from modulatory centers including histaminergic fibers from hypothalamus and serotonergic fibers from the dorsal raphe. Thus the retina provides a second example of complex sensory transformations carried out with little feedback from higher brain centers.

The finding that feedback was not necessary for the transformation of the bulb input into a concentration invariant output suggests the possibility that other relatively short term olfactory transformations such as adaptation and configural odorant responses (Kay et al., 2005; Coureaud et al., 2008) might also be carried out in the olfactory bulb without feedback from higher brain centers.

Determining the synaptic processes responsible for transforming the bulb input into a relatively concentration invariant output is simplified by the finding that it occurs in the olfactory bulb itself independently of feedback from higher brain centers.

## ACKNOWLEDGEMENT

We thank J.H. Choi and members of her laboratory at the Korea Institute of Science and Technology for generous help in making the recordings of the γ oscillations.

